# Enhanced production of nitrogenase components in *Nicotiana benthamiana* through co-expression with Bacterioferritin A

**DOI:** 10.64898/2026.06.29.734789

**Authors:** Alejandro M. Armas, Viviana Escudero, Julia Quintana, Mario Rodríguez-Simón, Isidro Abreu, Juan Andrés Collantes-García, Bidya B. Gupta, Elisa de Ansorena, Daniel Raimunda, Luis M. Rubio, Manuel González-Guerrero

## Abstract

- Engineering nitrogen fixing crops requires not only transferring the nitrogenase structural genes, but also the accessory genes to synthesize its iron-sulphur cofactors. Scaffold protein NifU is a critical element in this system as the starting point of nitrogenase cofactor assembly. NifU has been successfully produced in plants, however, its optimal production required high levels of iron in the medium. This is likely due to a faulty connection with the endogenous iron trafficking network
- To identify specific elements targeting iron to NifU, pull-down assays were performed to identify showing bacterioferritin A (BfrA) as a likely candidate. Co-immunopurification, mutant characterization, iron transfer assays, and co-expression in *Nicotiana benthamiana* assays were carried out.
- BfrA transfers iron to NifU through protein-protein interactions. When these two proteins were co-expressed in *N. benthamiana* leaves, there was an increase in NifU production. In turn, it led to doubling NifH synthesis, a nitrogenase structural protein that is also required for the synthesis of the more complex nitrogenase cofactors.
- Our results provide a new element towards engineering nitrogen-fixing crops. They also underscore the importance of transferring the metal delivery systems when expressing metalloproteins in heterologous systems.

## Introduction

Nitrogen is an essential and limiting nutrient in the biosphere (Smil, 1999b). Its most abundant source, N2, is only accessible to a few bacteria, archaea, and to organelle-like endosymbionts present in some unicellular algae. They achieve this by synthesizing nitrogenases, the only known enzymes that convert, fix, N2 into ammonia (Dos Santos *et al*., 2012). Towards ensuring an appropriate nitrogen supply, plants have established different symbiotic relationships with diazotrophic organisms, ranging from loose associations in the rhizosphere to endosymbiosis in specifically developed organs (Masson-Boivin *et al*., 2009) or even specialized organelle (Coale *et al*., 2024). However, biological nitrogen fixation is not sufficient to sustain modern agriculture and the demand from the current world population (Smil, 1999a, 1999b). It is estimated that over 50% of the nitrogen content of humans originates from nitrogen fertilizers produced through the Haber-Bosch process. This has had a huge economic and environmental costs due to the large energy requirements of the Haber-Bosch reaction and the polluting effect of run-off fertilizers and their derivates (Sutton *et al*., 2011). To ameliorate these problems, new ways to optimize nitrogen fertilization and nutrition of crops are being explored (Curatti & Rubio, 2014; Bloch *et al.,* 2020; Mus *et al*., 2016; Guo *et al*., 2023).

One of these approaches is engineering plants that can synthesize nitrogenase and directly use N2. These plants would express nitrogen fixation (*nif*) genes, and the resulting proteins are targeted to organelle, particularly mitochondria. There, Nif proteins will be protected from damaging O2 (López-Torrejón *et al*., 2016; Buren *et al.,* 2017; Johnston *et al.,* 2020; López-Torrejón *et al.,* 2021). However, these plants do not only need to express the nitrogenase structural proteins (NifH, NifD and NifK in nitrogenase most common form), but also the proteins synthesizing its three different iron-sulphur ([FeS]) cluster cofactors (Burén *et al*., 2020; Guo *et al*., 2023). Nitrogenase requires a [Fe4S4] group in each NifH2 dimer, and two P-clusters (Fe8S7) and two iron-molybdenum cofactors (FeMo-co; Fe7S9MoC-homocitrate) in each NifD2K2 tetramer. In diazotrophic organisms, the latter two complex clusters are synthesized from [Fe4S4] precursors made on scaffold protein NifU with sulphur provided by cysteine desulfurase NifS and an unknown iron donor (Dos Santos *et al*., 2004; Burén *et al*., 2020). Interestingly, eukaryotes are also able to synthesize [Fe4S4] groups in their mitochondria (Lill & Freibert, 2020). However, there is no proper connection between the endogenous [FeS] synthesis machinery and these heterologous Nif proteins, or the amount of [Fe4S4] synthesized is not sufficient for Nif proteins assembly. As a result, co-expression with NifU is often required to produce other Nif proteins in eukaryotes (Buren *et al*., 2019; Eseverri *et al*., 2020).

Beyond [FeS] cluster trafficking in mitochondria, iron availability itself is a limiting factor in nitrogenase synthesis (Miller and Orme-Johnson, 1992). Nitrogenase requires a total of 38 iron atoms in cofactors that are synthesized by ferro-enzymes (Burén *et al*., 2020). As a result, diazotrophic organisms and their hosts upregulate their iron uptake mechanisms to ensure a steady supply of this nutrient (Terry *et al*., 1991; Küpper *et al.,* 2008; Rosa-Núñez *et al*., 2023; Guío *et al.,* 2025). Considering the widespread iron deficiency in large areas of the world that already limits crop production (White & Brown, 2010), engineering nitrogenase in plants will also entail reinforcing iron uptake, as evidenced by the positive effect of iron-fertilization on NifU production in plants (Jiang *et al*., 2021). However, iron fertilization towards ensuring nitrogen fixation in nitrogenase-expressing crops is not a sustainable strategy when these technologies are implemented in the fields. Alternatively, the upregulation of iron uptake genes, as those used to iron-fortify plants (Vasconcelos *et al*., 2017), can ensure increased iron levels. However, this approach risks flooding the cells with iron that can become toxic at high levels or deplete even further the soils from this nutrient. To avoid these shortcomings, precision iron allocation to NifU could be used to maximize [Fe4S4] synthesis and transfer to other Nif components in plants.

To identify proteins transferring iron to NifU, we carried out pull-down assays using *A. vinelandii* protein extracts and found interaction with bacterioferritin A (BfrA, Avin14050), which was then shown to act as an iron donor to NifU. The co-expression of BfrA with NifU and NifH in *Nicotiana benthamiana* plants results in doubling the synthesis of both Nif proteins. These results show that transferring the endogenous iron transfer network proteins enhances nitrogenase [Fe4S4] cluster synthesis in plant mitochondria.

## Materials and Methods

### Biological material and growth conditions

*Azotobacter vinelandii* DJ was used as parental strain, *A. vinelandii* DJ984 containing a kanamycin resistance cassette integrated in *bfrA* was kindly provided by Prof. Dennis Dean (Virginia Tech, USA). This mutant strain was used to generate *A. vinelandii* DR1 by introducing a 2.1 kb DNA fragment amplified with the indicated primers (Table S1) containing the *bfrA* gene and the 800 bp flanking regions. This amplicon (400 ng) together with 10 ng of vector pDB303 were introduced in *A. vinelandii* DJ984. Allele exchange screening was performed by congression on BN media (Dos Santos, 2019) to identify rifampicin resistant and kanamycin sensitive strains. Allele exchange was confirmed by PCR and sequencing. *Escherichia coli* BL21 (DE3) pLysS was used to produce NifU and BfrA in LB medium. *Agrobacterium tumefaciens* GV3101 was used for plant transformation.

*Nicotiana benthamiana* plants were used in all transient expression experiments. Seeds were germinated directly on soil under high moisture conditions by covering trays with a plastic cover. Ten-day-old seedlings were transferred to 13 cm pots and grown for an additional 15 days prior to infiltration. Plants were grown in the greenhouse under long day conditions (16 hours natural light supplemented with artificial light for cycle completion / 8 hours dark). Plants were watered by irrigation (2.5 cm water level) once or twice a week with tap water.

### Protein purifications from *E. coli*

To produce TSBfrA in *E. coli* BL21 (DE3) pLysS, *bfrA* was PCR amplified with the primers listed in Table S1, that added *Nde*I and *BamH*I restriction sites to clone in pET16b(Strep). These cells were grown in LB media at 30°C until OD600 ∼0.6. Protein synthesis was induced with 0.5 mM IPTG, and cells were grown for 6 hours before being pelleted. Pellets were resuspended in lysis buffer A (50 mM Tris-HCl pH 8.0, 100 mM NaCl, 10% glycerol, and 1 mM phenylmethylsulfonyl fluoride (PMSF)). Cells were lysed in a French Press cell at 1,500 lb per square inch and cleared by centrifugation at 25,000 *x g* for 30 min at 4°C and filtration with 0.45 µm pore size syringe filters (Sartorius, Germany). Protein purification was performed at room temperature using 1 ml of Strep Tactin XT superflow resin high capacity (IBA Lifesciences, Germany) according to the manufacturer’s protocol. Briefly, cell free extracts were loaded onto the resin, then washed five times with 2 column volumes (CV) of buffer A. Protein of interest was then eluted with 50 mM biotin in buffer A in three steps (using 0.5, 2 and 2 CV respectively). Columns were regenerated following manufacturer instructions. Small molecules and salts were removed by dilution and concentration with centrifugal membrane devices (Amicon Ultra, Millipore, Burlington, MA, USA) following manufactureŕs recommendations.

The plasmid producing HBfrA was obtained by PCR using the primers listed in Table S1. HBfrA was cloned in a pET16b(His) vector previously digested with *Nde*I and *Bam*HI enzymes. *E. coli* BL21 (DE3) pLysS carrying the plasmid was grown in LB until OD 0.6 and induced with 0.5 mM IPTG, supplemented with 2 mM Cys and 0.2 mM ammonium Fe(III) citrate, for 6 h, 105 rpm, 30 °C. 10 g of the cells were resuspended in 30 ml buffer A (100 mM Tris pH 7.8, 250 mM NaCl), DNAse 5 μg/ml, 1 mM PMSF, lysed in a French Press and cleared by centrifugation. The protein was purified from the supernatant using a 5 ml Ni-NTA Agarose (QIAGEN, Germany) column, washed with 20 CV of 20 mM imidazole in buffer A, 20 CV of 40 mM imidazole in buffer A, and eluted with 20 ml 400 mM imidazole in Buffer A. The protein was desalted as above, and stored at -80°C in buffer A.

_H_NifU and NifUS were expressed and purified as indicated (Barahona *et al*., 2024).

All protein purification were performed within a COY chamber in anaerobic conditions (<5 ppm O_2_) (COY labs, USA), with all buffers degassed overnight.

### Pull-down assays and proteomics

Purified NifUS was used as bait to retain interacting proteins from cell-free extracts of diazotrophically grown *A. vinelandii* DJ, as previously described (Jimenez-Vicente *et al*., 2018). Briefly, *A. vinelandii* cells were pelleted, resuspended in buffer A, lysed, and cleared as indicated above for *E. coli*. 15 nmols of NifUS were loaded onto a 0.2 ml Strep-Tactin Superflow resin column (IBA Lifesciences), previously equilibrated with buffer A. The column was then washed 5 times with 2 CV of buffer A per wash to remove unbound bait protein. Then, 3 ml of 5 mg/ml *A. vinelandii* cell free extracts were loaded onto the column and washed 5 times with 2 CV of buffer A. Protein elution was carried out with 2.5 mM desthiobiotin in buffer A (3 steps with 2 CV each). Elution fractions were pooled and interfering molecules such as desthiobiotin removed by ultrafiltration using an Amicon® Ultra Centrifugal Filter, 3 kDa MWCO (Merck, USA). Eluted proteins were sent to LC-MS/MS analyses at the Unidad de Proteómica (Universidad Complutense de Madrid, Spain).

### Iron loading of BfrA

To load iron into BfrA, 2.5 ml of 12 μM BfrA was prepared in buffer P (100 mM KH2PO4 pH 7.6, 10 % glycerol, 20 mM ascorbate, 5 mM 1,4-dithiothreitol (DTT)) and incubated at room temperature for 40 min. DTT was eliminated with a PD10 column (GE Healthcare). Iron was added as (NH_4_)_2_Fe(SO_4_)2 in 10 steps with 15 min incubation in between until reaching the final concentration of 1.5 mM iron. Samples were incubated for 16 h at room temperature. The reconstitution mixture was diluted and concentrated using 30 kDa cutoff pore size centrifugal membrane devices (Merck) to remove excess iron and changing to buffer W (100 mM Tris pH 8, 150 mM NaCl).

### In vitro [Fe-S] cluster reconstitution

Purified tagged NifU variants were reconstituted *in vitro* as described (Barahona *et al*., 2024). Briefly, 20 μM of NifU dimer in 100 mM MOPS (pH 7.5) buffer, 9 mM 1,4-dithiothreitol (DTT) was incubated at 25°C for 30 min. 225 nM NifS, 1 mM L-cysteine and 0.4 mM (NH4)2Fe(SO4)2 were added. Iron and cysteine additions were divided in 10 steps of 15 min each until reaching the final concentration. The reconstitution mixture was kept in ice for 4 h and then desalted using 30-kDa cutoff pore size centrifugal membrane devices (Merck) to remove excess reagents and exchange to Buffer W. Reconstituted NifU proteins were stored in liquid N2 until use.

### Iron transfer assays

Iron transfer assays were carried out for 5 min using 50 μM of NifU monomer and 50 μM of iron-loaded BfrA monomer, 0.8 mM ethylene glycol-bis(β-aminoethyl ether)-N,N,N′,N′-tetraacetic acid (EGTA), 2.5 mM Tris(2-carboxyethyl)phosphine (TCEP) in 200 μl under anaerobic conditions. The solution was passed 3 times through a 200 μl Strep-Tactin XT 4Flow high-capacity resin (IBA Lifesciences), previously equilibrated with anaerobic buffer W. This flowthrough was loaded onto a 1 ml Sephadex G-25 column previously equilibrated with anaerobic buffer W to remove unbound iron. Protein and iron content in the collected fractions were determined by the BCA method (Pierce, USA) and by Atomic Absorption Spectroscopy, respectively.

To control for the transfer of possible released free iron, HNifU and TSBfrA were incubated for 5 min separated by a 2-kDa pore-size cutoff dialysis membrane, previously equilibrated for 1 h with buffer W. Samples from both membrane sides were collected to determine the protein and iron concentration.

### Nitrogenase activity

The acetylene reduction assay was used to determine nitrogenase activity. When assessing this activity in cell, acid-washed flasks were used to grow 250 ml of *A. vinelandii* for 16 h in Burk media. The following day, after refreshing the cultures in the same media for 2 h, they were pelleted for 10 min at 1,500 *x g* and resuspended in Burk media without combined nitrogen (diazotrophic conditions) with and without FeCl3 for 4 h. Low-iron conditions were produced by using a Burk media with no added iron. One ml of culture was incubated with 0.5 ml of acetylene (6 %) at 30 °C for 15 min in hermetically closed vials. After stopping the reaction with 0.1 ml of 8 M NaOH, a 50 μl sample was analysed in a gas chromatograph (Shimadzu GC-8A, Japan). The quantity of ethylene produced was standardized by the OD600 of each culture.

*In vitro* reconstitution of apo-NifH by NifU was determined as described (Shah *et al*., 1986; Jiang *et al.,* 2021). Reactions were prepared inside anaerobic chambers. 4 μM of apo-NifH was incubated with 0.1 μM of NifDK, 40 μM NifU and ATP-regenerating mixture (1.23 mM ATP, 18 mM phosphocreatine, 2.2 mM MgCl_2_, 3 mM DTH, 5 mM DTT, 46 μg/ml of creatine phosphokinase, 100 mM MOPS pH 7.4) in a final volume of 600 μl inside 9 ml serum vials under argon atmosphere containing 500 μl acetylene (1 atm). The reaction was carried out at 30 °C in a shaking water bath for 15 min. Reactions were stopped by adding 100 μl of 8 M NaOH. Ethylene formed was measured in 50 μl gas phase samples as previously indicated.

### Confocal imaging

The *bfrA* DNA sequence was synthesized codon-optimized for *N. benthamiana* and made compatible for MoClo cloning (Genscript, USA) (Weber *et al*., 2011; Werner *et al*., 2012). (Table S2). Modular pieces were used to develop a transcriptional unit containing the 35S promoter, SU9 mitochondrial signal peptide, *egfp*, *bfrA*, and the 35S terminator region (Table S2) (Eseverri *et al*., 2020; Jiang *et al*., 2021). This module was cloned in pICH47732 and introduced in *Agrobacterium tumefaciens* GV3101 to transform *N. benthamiana* leaves. mitoRFP was co-agroinfiltrated to serve as mitochondria locating signal (Candat *et al*., 2014). Agrobacterium feeder cultures containing the gene of interest in a binary vector (see primers and constructs) were used to inoculate liquid LB with rifampicin 25 μg/ml, gentamycin 100 μg/ml and the corresponding binary vector resistance to a final dilution 1:100. Cultures were grown at 28 °C for 16–24 h to the stationary phase. Cells were collected by centrifugation at 3,500 *x g* 15 min at 10 °C in a swinging bucket rotor and gently resuspended in freshly prepared infiltration solution (10 mM MgSO_4_, 10 mM MES pH 5.5 and 150 μM acetosyringone) to a final cumulative OD of 0.9. Cultures were incubated for 3 h in darkness to induce virulence and used for to infiltrate 1 ml of solution per leaf into leaves 3 and 4 of *N. benthamiana* plants on three-week-old plants.

Leaf discs were obtained 4 days-post-inoculation (dpi) and observed in a Zeiss LSM 880 laser scanning confocal microscope, using excitation wavelengths of 488 and 561 nm for GFP and mitoRFP observation, respectively.

### Transient expression in *N. benthamiana* and protein purification

A twin-strep tag was used to purify *N. benthamiana* codon-optimed *A. vinelandii* BfrA, *A. vinelandii* NifU, and *H. thermophilus* NifH. Transcriptional units were assembled by MoClo and finally cloned in level 2 acceptor vector pAGM4673 (Weber *et al*., 2011; Werner *et al*., 2012) (Table S2). The BfrA module consisting of AtUbq promoter10:SU9:BfrA:35S terminator was cloned by In-fusion (Takara Bio, Japan) by amplifying the transcriptional unit using primers 5*BamH*IBfrAInf and 3*BamH*IBfrAInf (Table S1) and cloning into the L2 module containing the transcriptional unit for TSNifH. NifH, NifM, p19, and NifU transcriptional units were produced in (Jiang *et al*., 2021).

Agroinfiltrated 4 dpi leaves (100-200 g) were collected for protein purifications. These were carried out in a glove box using degasified buffers as indicated (Jiang *et al*., 2021). Briefly, homogenize 1:1 (weight:volume) in lysis buffer (0.1 M Tris HCl pH 8.6, 0.2 M NaCl, 5 μg/ml DNaseI, 1 μg/ml leupeptin, 1 mM PMSF, and 5 mM β-mercaptoenanol) using a blender for 5 min. Samples were filtered through miracloth and centrifuged at 54,000 x *g* for 1 h at 4 °C. Filter supernatant through a 0.2 μm filter and pass through a Strep-Tactin XT 4Flow high-capacity resin (IBA Lifesciences) equilibrated with buffer W. Wash with 10 CV of Buffer W and elute with 5 CV of 1.2 % biotin in buffer W. Samples were concentrated and biotin diluted out by successive dilutions. Proteins were frozen and stored under anaerobic condition in 20% glycerol in buffer W.

### Iron quantification

Protein samples were mineralized in 37.5 % analytic grade nitric acid for 10 min at 80 °C. Samples were then diluted to a total concentration of 2 % nitric acid with ultrapure water. To determine iron content in cellular samples, 1 ml of the cultures was pelleted and digested in 50 μl 6 N HCl overnight. Samples were diluted 10-fold to reduce acid concentrations. Iron concentrations were determined in an Atomic Absorption Spectrometer ContrAA 800G (AnalytikJena, Germany) using commercially available analytic grade metal standards (Inorganic Ventures, USA).

## Results

### *Azotobacter vinelandii* NifU interacts with BfrA

To identify putative iron donors to NifU, a C-terminal Twin-Strep-tagged NifU (NifUTS) was used as bait to pull down proteins produced by *A. vinelandii* wild type (DJ strain) 4 hours after being transferred to diazotrophic growth conditions. Among the 20 most abundant proteins found to co-elute with NifUTS (Table 1) were the previously reported cysteine desulfurase NifS, which provides sulfur for Fe-S cluster assembly on NifU (Smith *et al*., 2005), and the molybdenum donor NifQ, which obtains its [Fe4S4] cluster from NifU (Barahona *et al*., 2024). While some of the remaining proteins isolated in the pull-down assay were known biotinylated proteins (such as pyruvate carboxylase), another, BfrA, had been characterized to be involved in iron homeostasis. This protein is a 24-mer protein involved in iron storage and remobilization (Rivera, 2017).

**Table 1.** Twenty most highly detected proteins in the pull-down assays with NifUS.

| Accession | Description | Coverage [%] | # Peptides | # PSMs | # Unique Peptides |
| --- | --- | --- | --- | --- | --- |
| C1DH18 | Nitrogen fixation protein NifU | 88 | 27 | 351 | 27 |
| M9YD16 | Cysteine desulfurase NifS | 27 | 8 | 50 | 8 |
| C1DH60 | Pyruvate carboxylase subunit B | 24 | 13 | 28 | 13 |
| P22759 | Bacterioferritin A | 46 | 8 | 21 | 8 |
| M9YB39 | Chaperone protein DnaK | 21 | 13 | 19 | 12 |
| M9YHW8 | 50S ribosomal protein L2 | 46 | 11 | 16 | 11 |
| M9YJZ7 | Glyceraldehyde-3-phosphate dehydrogenase | 31 | 14 | 16 | 2 |
| M9YMW1 | NAD-dependent aldehyde dehydrogenase | 29 | 13 | 15 | 1 |
| C1DH59 | Pyruvate carboxylase subunit A | 25 | 11 | 13 | 11 |
| M9YH82 | DNA topoisomerase 4 subunit A | 11 | 8 | 9 | 8 |
| C1DEN2 | Glutamate--tRNA ligase | 15 | 8 | 9 | 8 |
| M9YIC3 | Translation initiation factor IF-2 | 11 | 8 | 8 | 8 |
| M9YAD8 | 2-oxoglutarate dehydrogenase E1 component | 8 | 7 | 7 | 7 |
| C1DKN7 | 30S ribosomal protein S4 | 27 | 6 | 7 | 6 |
| P11068 | Molybdenum donor NifQ | 32 | 7 | 7 | 7 |
| M9YE09 | 50S ribosomal protein L21 | 48 | 5 | 6 | 5 |
| C1DGZ8 | Nitrogenase molybdenum-iron protein NifK | 12 | 6 | 6 | 6 |
| M9YK53 | 50S ribosomal protein L24 | 37 | 4 | 5 | 4 |
| C1DQ78 | 50S ribosomal protein L13 | 32 | 4 | 5 | 4 |
| C1DFK5 | Polyribonucleotide nucleotidyltransferase | 9 | 5 | 5 | 5 |

N-terminal Twin-Strep tagged BfrA (TSBfrA) was purified from recombinant *E. coli* cells (Fig. S1) containing around 1 iron atom per BfrA monomer (Table 2). BfrA could be loaded *in vitro* with additional iron to average ∼15 iron per monomer (Table 2), indicating that the 24-mer cage is formed (∼ 360 iron atoms per cage). Both as-isolated and reconstituted (R) iron-loaded TSBfrA were incubated with N-terminal His6-tagged NifU (HNifU) to validate their interaction through co-purification assays. HNifU co-purified with TSBfrA only when the latter was loaded with iron, while no interaction could be observed with as-isolated BfrA (Fig. 1, Fig. S2). This was regardless of the iron levels of HNifU, since both the as-isolated form with 2 iron per monomer and the form in which the [Fe_4_S4] clusters were reconstituted (R-HNifU) (Table 2) showed the same pattern of interaction with BfrA (Fig. 1a-d). However, the incubation with the as-isolated HNifU seemed to be the strongest one (Fig. 1a). This interaction was also validated when the tags were switched and iron-loaded N-His9 tagged BfrA (HBfrA) was incubated with as-isolated C-Strep tagged NifU (NifU_S_) (Fig. S3). To determine the role of the transient [FeS] coordinating sites in R- TSBfrA interaction with HNifU, Cys residues 35, 62, 106, 272, and 275 were replaced by Ala to produce CA-HNifU (Smith *et al*., 2005; Collantes-García *et al*., 2026) (Table 2). In this situation, no interaction could be observed, even when the core [Fe_2_S_2_] cluster was removed from CA-HNifU (Fig. 1e,f).

**Figure 1.**
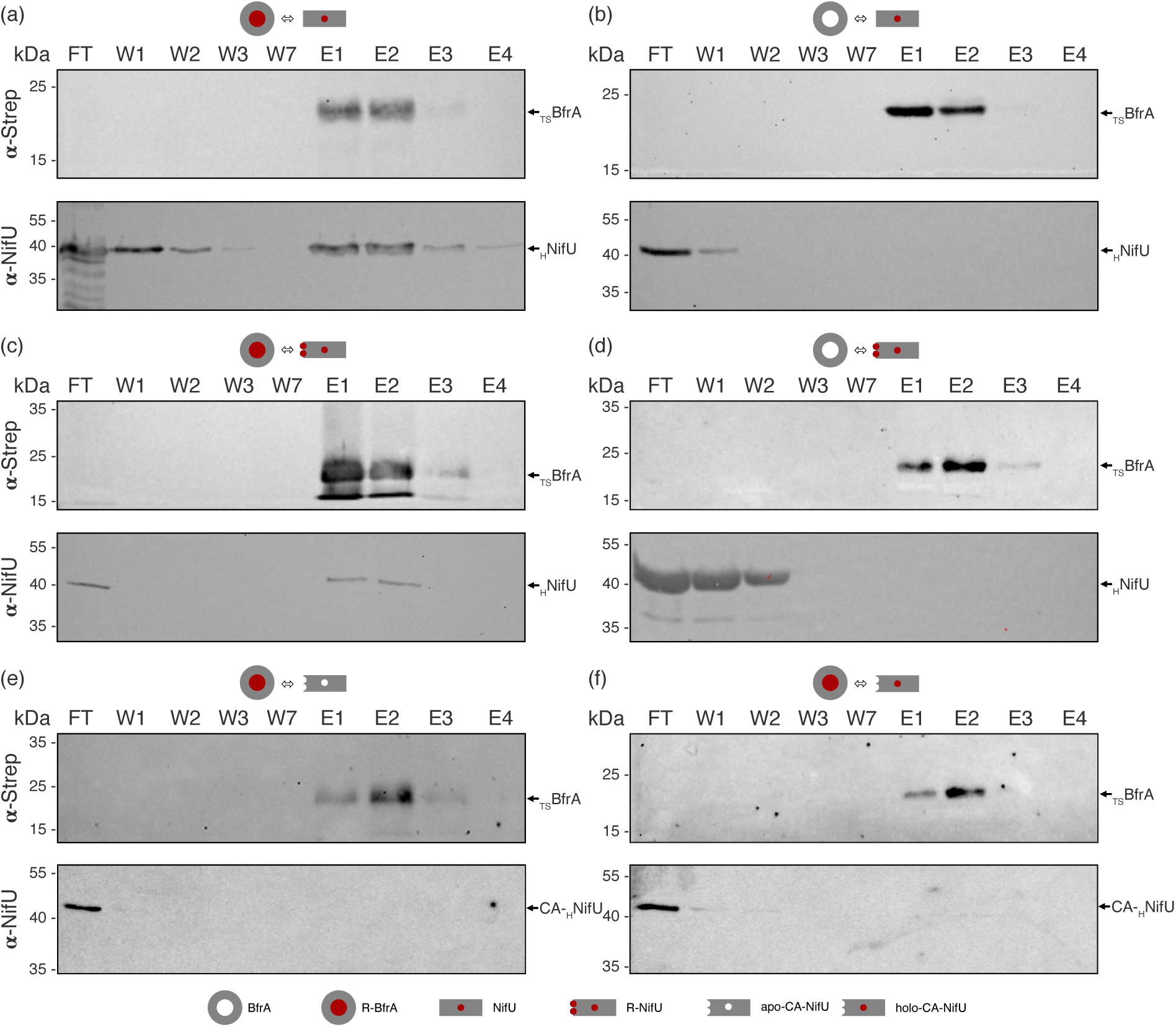
Iron-loaded BfrA interacts with active NifU. Top panel shows the immunodetection of Strep-tagged iron-loaded (a, c, e) or as-isolated (b, d, f) BfrA using an anti-Strep antibody in flowthough (FT), washes (W1, W3, and W7), and elution (E1, E2, E3, and E4) fractions after being incubated with a histidine-tagged as-isolated (a,b), or reconstituted (c,d) wild type NifU or with apo or holo-CA-NifU (e, f), and passed through a Strep-column. Bottom panel shows the immunoblot of the same fractions developed with an anti-NifU antibody. Images show a representative assay (n=3). Full length immunoblots are shown in Supplementary Figure S2.

**Table 2.**
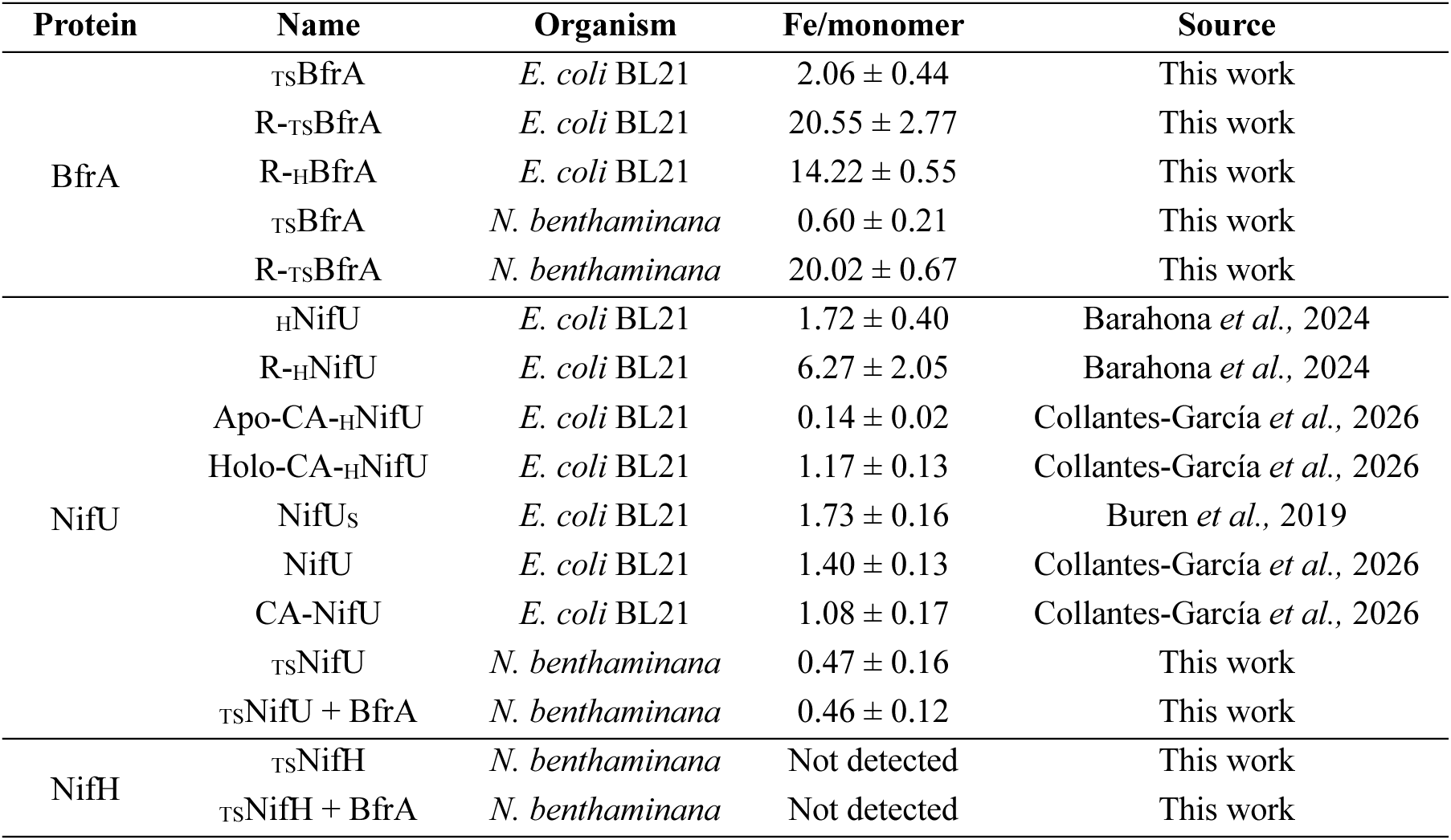
Proteins purified in this study and their iron content (Mean ± SD, n=3).

### BfrA donates iron to NifU through protein-protein interactions

The interaction between NifU and BfrA and its dependency on iron is indicative of a role of BfrA in providing this cation to NifU to initiate nitrogenase cofactor assembly. In this model, mutation of *bfrA* would lead to growth defects and a reduced nitrogenase activity because of lower cofactor assembly. To test this hypothesis, a *bfrA* insertional mutant was produced and grown under diazotrophic and non-diazotrophic conditions (Fig. 2). Under non-diazotrophic conditions, no large differences on growth were observed under iron replete conditions, although significant differences could be observed when grown in a low-iron medium (Fig. 2 a,b). These differences between wild type and *bfrA* strains were larger under diazotrophic conditions (Fig. 2 c,d). In this scenario, *bfrA* grew worse both in an iron sufficient and in an iron deficient environment. In all scenarios, the phenotype reverted to wild-type levels by reintroducing a wild-type *bfrA* gene in the *bfrA* strain. The reduced growth under diazotrophic conditions was the likely consequence of the lower iron content (Fig. 2e) and nitrogenase activity (Fig. 2f) of the *bfrA* strain compared to the wild type and the *bfrA* complemented strains.

**Figure 2.**
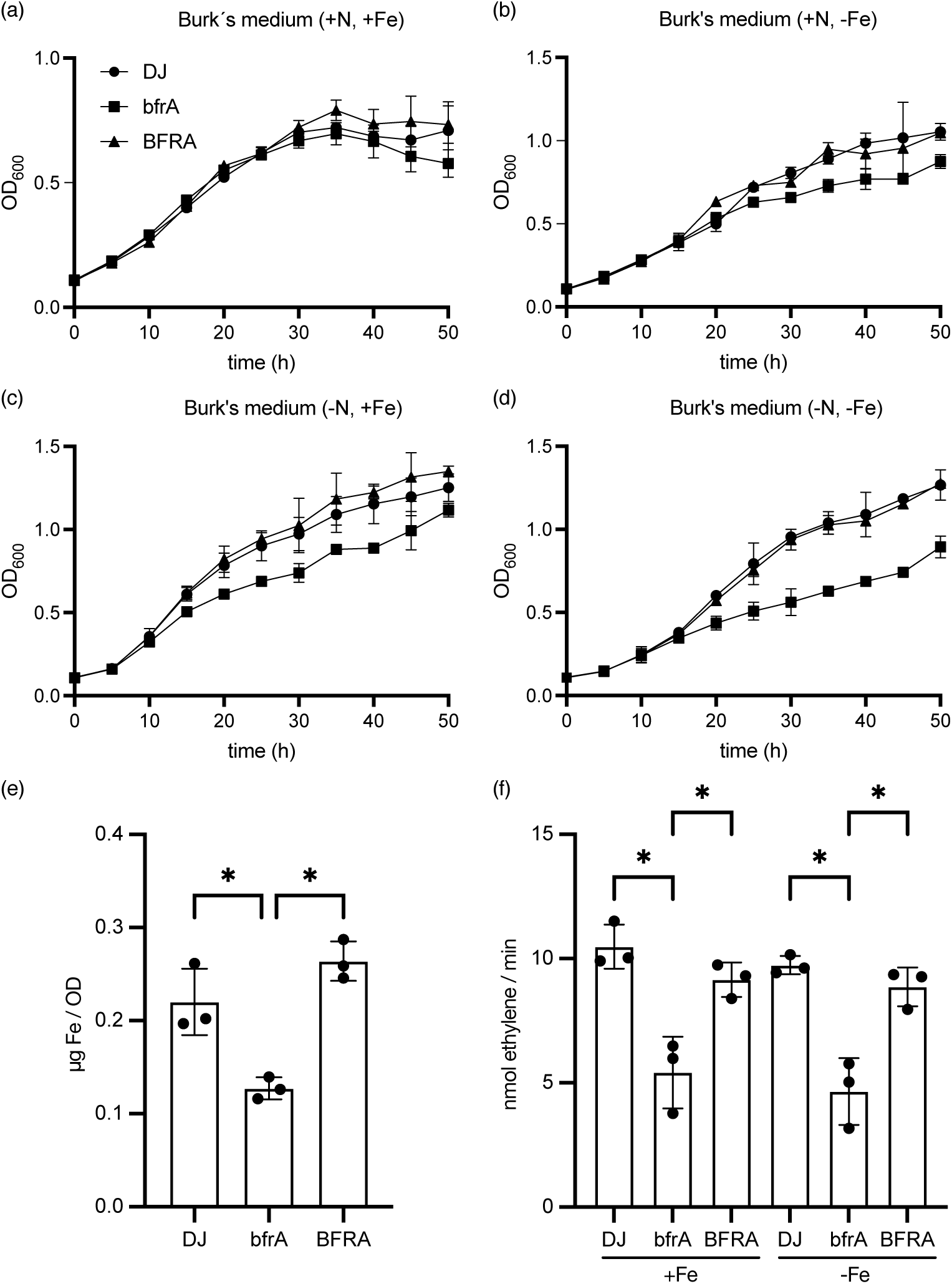
BfrA is required for optimal nitrogen fixation in *A. vinelandii*. a) Growth under non-diazotrophic conditions of wild type *A. vinelandii* strain (DJ), *bfrA* insertional mutant (*bfrA*), and *bfrA* transformed with a wild-type copy of *bfrA* (BFRA) in Burk’s medium. b) Growth under non-diazotrophic conditions of *A. vinelandii* strains DJ, *bfrA*, and BFRA in iron-deficient Burk’s medium. c) Growth under diazotrophic conditions of *A. vinelandii* strains DJ, *bfrA*, and BFRA in Burk’s medium. d) Growth under diazotrophic conditions of *A. vinelandii* strains DJ, *bfrA*, and BFRA in iron-deficient Burk’s medium. e) Iron content in *A. vinelandii* strains DJ, *bfrA*, and BFRA grown in Burk’s medium in diazotrophic conditions. f) Nitrogenase activity of *A. vinelandii* strains DJ, *bfrA*, and BFRA grown in Burk’s medium in diazotrophic conditions 4 h after de-repression. Bars represent the average ± SD (n=3). * Indicates statistically significant difference (*p* <0.05).

To confirm a direct role of BfrA on transferring iron to NifU, first iron binding of free iron was tested using as-isolated HNifU, indicating that one iron atom could be bound in addition to the other two from the [Fe2S2] cluster in the core domain (Fig. 3a). To ascertain that BfrA transferred iron to NifU, R- _TS_BfrA was incubated with a tag-less NifU containing just the [Fe2-S2] of the core domain (Table 2). Since reducing agents promote iron release from bacterioferritins (Yao *et al.,* 2012), 2.5 mM TCEP was added to the buffer when transfer to NifU was assessed. 0.8 mM EGTA was added to bind labile iron and to restrict iron transfer to direct protein-protein interactions. Iron and protein content were determined in the flowthrough fraction were only NifU and no TSBfrA was released (Fig. 3b, Fig. S4). In this condition, a ratio of around three irons per NifU monomer could be detected, which was significantly different to the NifU approximate 1:1 ratio obtained when no R- TSBfrA was added to the buffer, or when no iron was released due to the absence of reducing agent (Fig. 3b). To further confirm that the iron detected in NifU resulted from the physical interaction with R- TSBfrA and not from iron dissociation and diffusion, similar determinations were made when the two proteins were separated by a dialysis membrane, resulting in no iron binding to NifU (Fig. 3b). Furthermore, no iron transfer was detected to CA-NifU (Fig. 3b).

**Figure 3.**
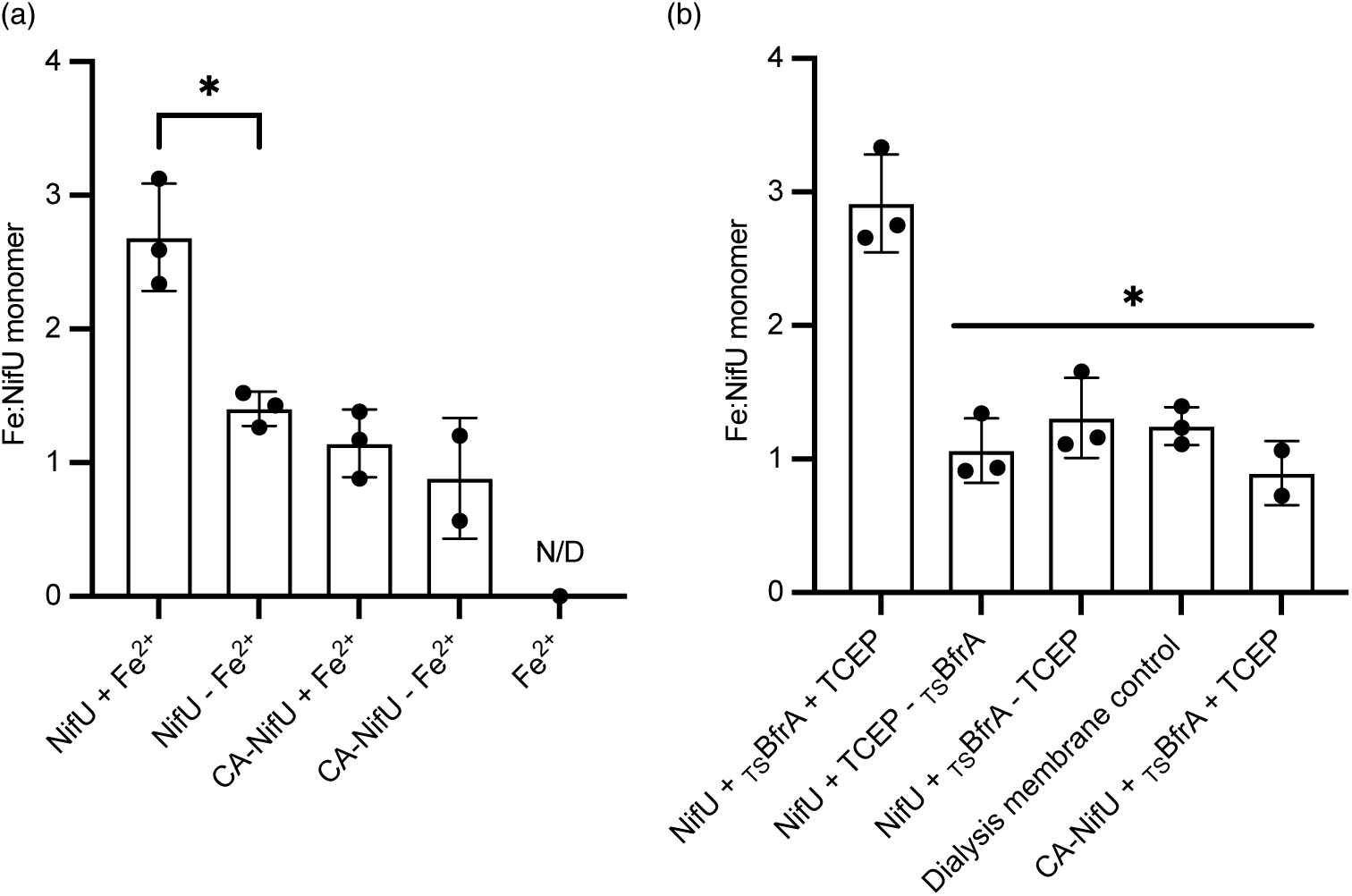
NifU obtains iron from BfrA. a) Iron content in elution fractions of HNifU or CA-HNifU as isolated (-Fe^2+^) or after incubation with a 10-molar ratio of Fe^2+^ (+Fe^2+^). The elutions were collected after passing the protein (+ Fe^2+^solutions) through Sephadex G-25 columns. A sample only with Fe^2+^ but no protein was used to control the removal of all the unbound Fe^2+^. b) Iron content in NifU or CA-NifU after incubation with iron-loaded TSBfrA and separation through a Strep-tactin column. In the dialysis membrane control, the two proteins were separated by a dialysis membrane that prevented protein diffusion but allowed for Fe^2+^ movement. Bars represent the average ± SD (n=3). * Indicates statistically significant difference (*p* <0.05).

### Co-expression with BfrA improves NifU and NifH synthesis in *Nicotiana benthamiana* leaves

To test whether the production of nitrogenase cofactor assembly proteins could be improved in plants by introducing iron transfer proteins, BfrA was codon-optimized to be expressed in *N. benthamiana* plants (Fig. S5). To target the protein to mitochondria, the SU9 signal peptide was fused to N-terminus of BfrA, with GFP to detect the protein with confocal imaging. The GFP signal colocalized with mitochondrial marker mitoRFP (Fig. 4a). No signal was observed when examining leaves not expressing GFP-BfrA (Fig. S6). TSBfrA was purified from *N. benthamiana* leaves and shown to be able to accumulate iron in reconstitution assays (Fig. S7; Table 2). To determine the effect of BfrA on NifU synthesis *in planta*, *N. benthamiana* leaves were infiltrated with a DNA fragment containing either N-terminal TS-tagged NifU (TSNifU), NifS and p19, or BfrA, TSNifU, NifS, and p19 (Fig. 4b). TSNifU was purified from both sets of plants with similar levels of purity (Fig. 4c). While the co-expression with BfrA did not significantly improve the iron levels of NifU (Table 2), the amount of NifU synthesized almost doubled (Fig. 4d). No additional increase in NifU synthesis was observed when the plants co-expressing *BfrA* where fertilized with iron.

**Figure 4.**
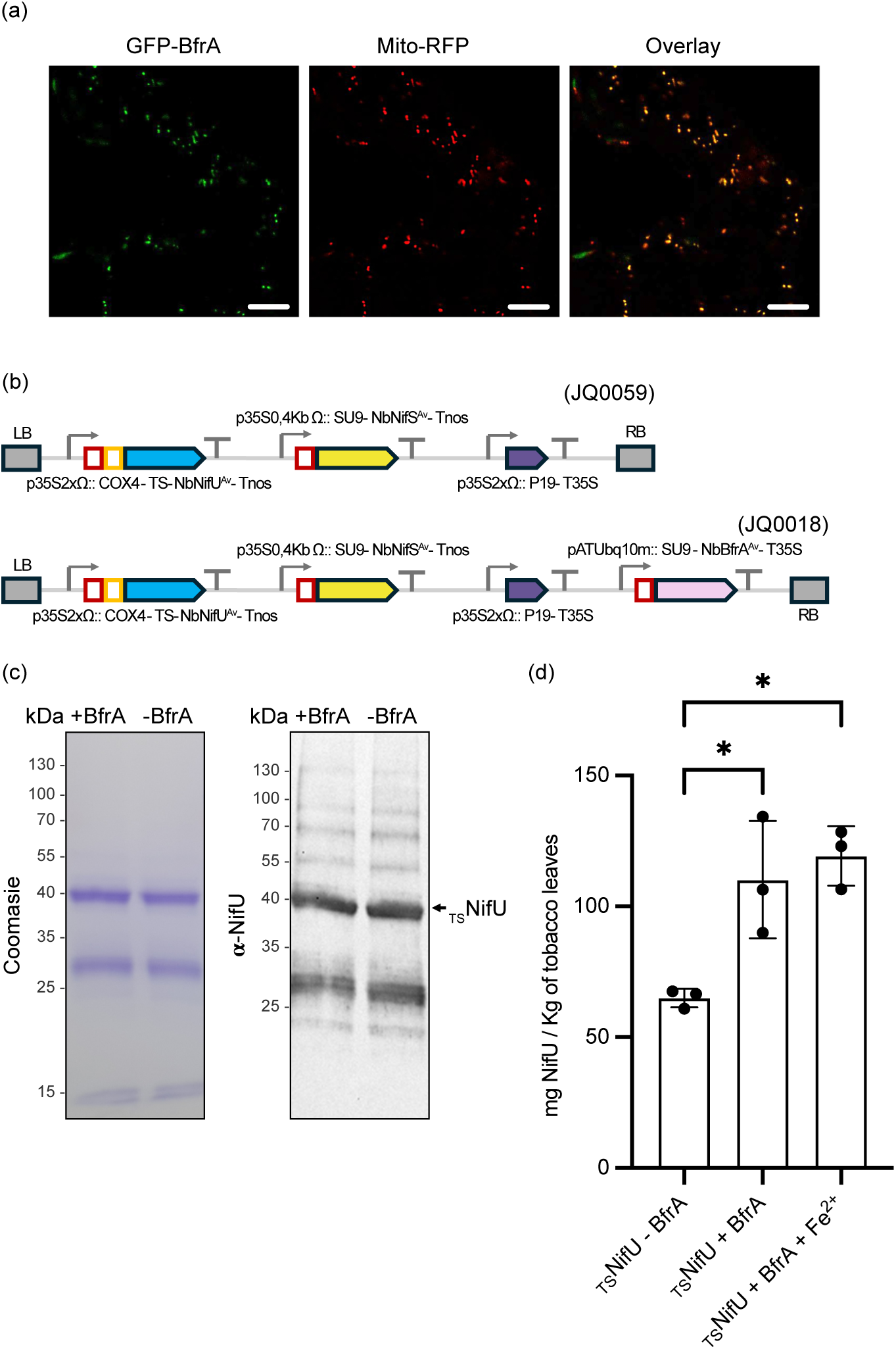
Mitochondria-targeted BfrA increases NifU yields in *N. benthamiana* leaves. a) Localization in *N. benthamiana* leaves of mitochondria-targeted GFP-tagged BfrA (left panel, green), mitochondria (centre panel, red), and the overlay of the two channels (right). Bars = 10 μm. b) Constructions used to produced TSNifU in *N. benthamiana* leaves in the presence (JQ0018) or absence (JQ0059) of BfrA. c) Commassie brilliant blue stain (left) and anti-NifU immunoblot (right) of TSNifU purified from *N. benthaminana* leaves co-expressed or not with BfrA. d) Purified TSNifU yields from *N. benthamiana* leaves when no BfrA was co-produced, when it was produced, and when the later plants were fertilized with iron. Bars represent the average ± SD (n=3). * Indicates statistically significant difference (*p* <0.05).

The transfer of [Fe4S4] groups from NifU to NifH is essential for NifH functioning and for the maturation of the other nitrogenase clusters. To test whether BfrA-based enhanced NifU synthesis resulted in increased NifH production/stability, N-terminal TS-tagged NifH (TSNifH) was co-expressed with NifS, p19, NifU, and NifM in *N. benthamiana* leaves in combination or not with BfrA (Fig. 5a). The NifH selected originated from *Hydrogenobacter thermophilus* as it was the NifH variant with best production in *N. benthamiana* (Jiang *et al*., 2021). All the other Nif proteins originated from *A. vinelandii.* TSNifH was purified from these plants to similar levels of purity (Fig. 5b). While the yields of TSNifH obtained from BfrA-synthesizing plants were 2.5x than those from the plants without BfrA (Fig. 5c), these proteins carried very little iron (Table 2). Consistent with this, no nitrogenase activity was observed when using as-isolated TSNifH as electron carrier to NifDK. However, when these proteins were reconstituted *in vitro* by providing the [Fe_4_S_4_] clusters, nitrogenase activity was recovered (Fig. S8).

**Figure 5.**
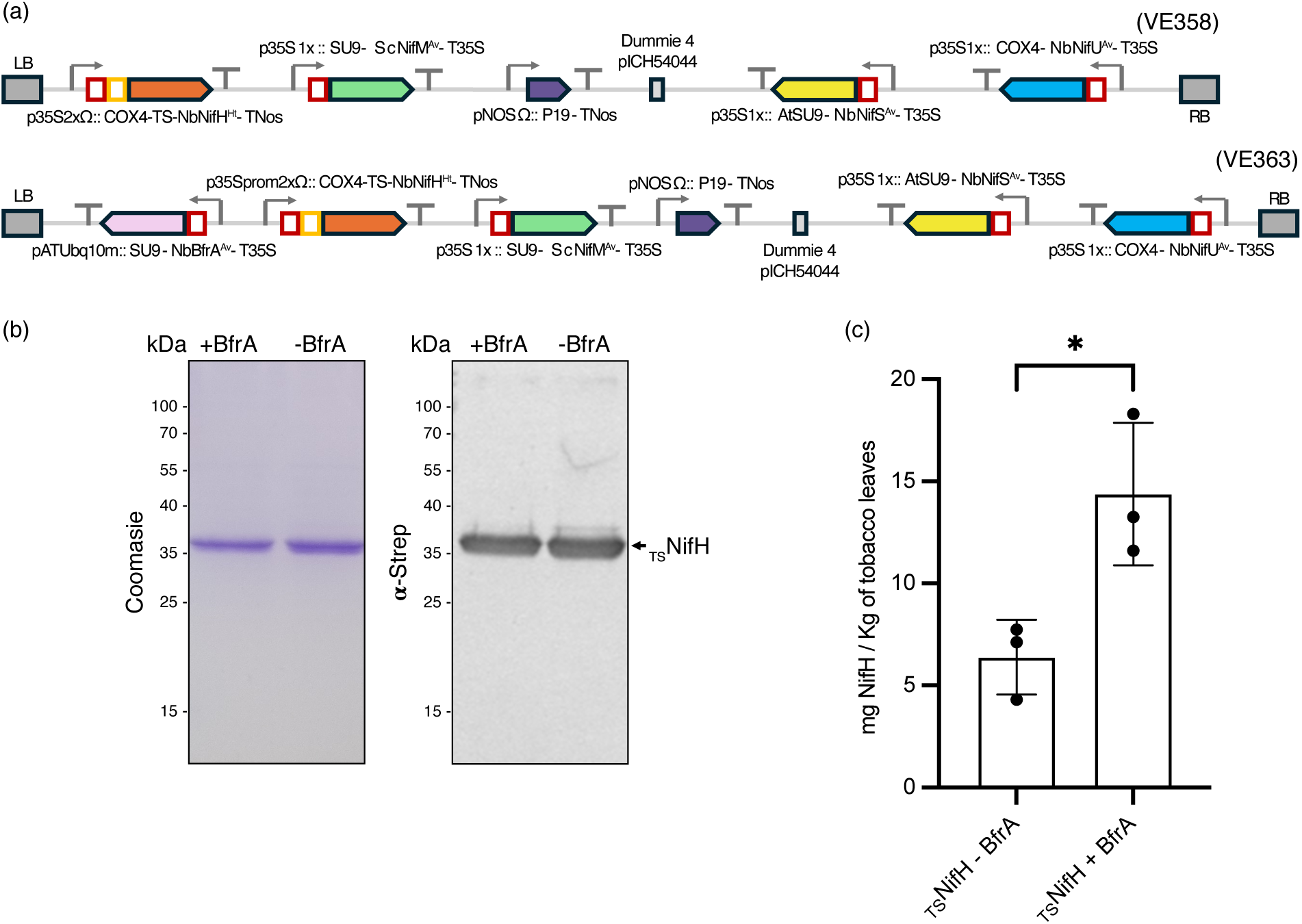
Mitochondria-targeted BfrA increases NifH yields in *N. benthamiana* leaves. a) Constructions used to produce TSNifH in *N. benthamiana* leaves in the presence (VE363) or absence (VE358) of BfrA. b) Commassie brilliant blue stain (left) and anti-NifH immunoblot (right) of TSNifH purified from *N. benthaminana* leaves co-expressed or not with BfrA. c) Purified TSNifH yields from *N. benthamiana* leaves when no BfrA was co-produced or when it was produced. Bars represent the average ± SD (n=3). * Indicates statistically significant difference (*p* <0.05).

## Discussion

Expressing functional nitrogenase in plants does not simply entail the synthesis of the apo-proteins, but also the production and insertion of the three different metalloclusters required for the catalysis (Burén & Rubio, 2018; Guo *et al*., 2023). This is a major bottleneck of engineering nitrogen fixing crops, as these co-factors are complex to synthesize and highly sensitive to oxygen. Furthermore, since iron is a growth-limiting nutrient in many soil types, plants have developed very precise mechanisms to control its use and allocation (Andresen *et al*., 2018). As a consequence, while nitrogenase components have been targeted to iron-rich organelle such as mitochondria and chloroplast, where the simpler [Fe_4_S_4_] cluster can be assembled, the production of many of the nitrogenase components still rely in co-expression with nitrogenase-specific [Fe4S4] scaffold protein NifU (Lopez-Torrejon *et al*., 2016; Buren *et al*., 2019; Eseverri *et al*., 2020; Jiang *et al*., 2021, 2022). These results indicate that expression of NifU is essential for nitrogenase cofactor synthesis in plants. However, NifU is still not fully “plugged-in” in the plant iron transfer network as evidenced by the requirement of iron fertilization to increase NifU yields (Jiang *et al*., 2021), indicating that an iron “overflow” would be needed for it to reach NifU. Introducing dedicated iron-delivery systems to NifU in the engineered plants might ameliorate this problem by ensuring a direct allocation of iron precursors.

Pull-down assays and co-purification experiments show that BfrA interacts with NifU. This interaction requires iron to be accumulated within the cage formed by the BfrA 24-mer, indicating a connection between iron storage and NifU activity. In *in vitro* assays, BfrA can act as iron donor to NifU, a process that requires protein-protein interactions, as shown by iron not being transferred when the two proteins are separated by a dialysis membrane. It is likely that the iron delivered to NifU is used to synthesize the [FeS] clusters that will be employed by nitrogenase. Supporting this hypothesis is the observation that the NifU cysteines used for [Fe4S4] cluster synthesis are required to accept iron from BfrA and even to establish a stable interaction between the two proteins. Furthermore, as expected for a protein involved in nitrogenase cofactor maturation, mutation of *BfrA* limits growth in diazotrophic conditions even during iron sufficiency. However, BfrA is also required for additional iron-dependent processes not linked to nitrogen fixation as evidenced by the growth reduction under non-diazotrophic growth in an iron-limiting situation. Similarly, BfrA would not be the only protein delivering iron to NifU since nitrogenase activity is not completely lost in *bfrA* mutants. However, this alternative system(s) would not likely include other (bacterio)ferritins in *A. vinelandii*, as none of these proteins co-eluted with NifU in the pull-down experiments.

Since BfrA is used for optimal nitrogen fixation rates in *A. vinelandii*, most likely by transferring iron to NifU, we hypothesized that it could be used to more efficiently allocate iron to plant-produced NifU. However, to test this possibility, iron content of the plant-purified NifU could not be used as evidence, since its [Fe4S4] clusters are extremely labile and likely lost during purification (Burén *et al*., 2020; Jiang *et al*., 2021, 2022; Baysal *et al*., 2022). Alternatively, increased NifU production was used as an indicator of iron reaching NifU and stabilizing it. This is based on the enhanced NifU synthesis observed in *N. benthamiana* upon iron fertilization (Jiang *et al*., 2021). Co-expression with *BfrA* results in higher NifU yields than when no BfrA is present, being consistent with iron-mediated NifU estabilization. The increased NifU production rate obtained is similar in that of iron fertilization (Jiang *et al*., 2021), but no synergy was observed by combining BfrA production and iron fertilization. This suggests that a saturation point has been reached. More importantly, the enhanced NifU production by BfrA also translates to NifH synthesis, an essential structural element of nitrogenase and a key element in FeMo-co maturation (Burén *et al.,* 2020). Since no interaction between BfrA and NifH was observed in the pull-down assays, this finding would be the consequence of the improved NifU levels. Since NifU synthesizes and transfers to NifH these clusters, what increases the stability of the protein, (Zhao et al., 2007), it can be hypothesized that the increased yields of NifH in *N. benthamiana* could follow a similar process. However, NifH was purified with no detectable iron, as already reported by other authors (Eseverri *et al*., 2020; Jiang *et al*., 2021; Baysal *et al*., 2022), indicating that either no iron was ever transferred in planta to NifH, or more likely, these clusters were lost during purification as damaging oxidative species are released during plant homogenization. Supporting this hypothesis is the observation that purified NifH was properly folded, since it could be reconstituted *in vitro*. Therefore, co-expression with BfrA will be a viable and more sustainable alternative to iron fertilization when engineered nitrogenase-producing crop are produced. Furthermore, these results are also proof-of-concept of the importance of identifying and co-expressing metal donors when engineering metal-dependent processes in plants.

## Acknowledgements

This work was by grant PID2021-12460OB-100 from the Ministerio de Ciencia, Innovación/Agencia Estatal de Investigación/10.13039/50110001103 and “ERDF A way of making Europe to MG-G. This work was supported by the Bill and Melinda Gates Foundation grant INV-005889 and the Gates Foundation grant INV-067006 to LMR. AMA was funded by a María Zambrano contract. JQ was supported by PCI2021-122052-2B from MICIN / AEI / 10.13039/501100011033 and by European Union NextGenerationEU/PRTR. IA was the recipient of a Juan de la Cierva-Formación postodoctoral fellowship from Ministerio de Ciencia, Innovación y Universidades (FJCI-2017-33222). JAC-G was supported by Formación de Personal de Investigación fellowships PRE2022-101253, funded by Ministerio de Ciencia, Innovación, y Universidades/Agencia Estatal de Investigación/10.13039/50110001103 and FSE+. The proteomic analysis was performed in the Proteomics Unit of Complutense University of Madrid, a member of ProteoRed and is supported by grant PT17/0019, of the PE I+D+i 2013- 2016, funded by ISCIII and ERDF.

## Competing interests

None declared

## Author contributions

AMA carried out the interaction assays between BfrA and NifU, determined the *bfrA* phenotype, and developed the iron transfer assay assisted by DR. VE performed the NifU and NifH purifications from *N. benthamiana* assisted by MR-S and BBG. JQ was responsible for targeting BfrA to *N. benthamiana* mitochondria and cloning the *bfrA* level 1 units. IA performed the initial pull-down assays. JAC-G determined NifH activities together with MR-S and EdA. LMR and MG-G obtained the funding, supervised the research, analysed results, and prepared the manuscript with input from all the authors

## Data availability

All the data used are included in this manuscript.

## Supporting information

**Figure S1. TSBfrA purification from *E. coli*.**

**Figure S2. Uncropped gels of Fig. 1**.

**Figure S3. NifUTS interacts with iron-loaded HBfrA.**

**Figure S4. Control for SBfrA contamination in the samples used for NifU iron content in Fig. 3b**.

**Figure S5. Sequence of *A. vinelandii BfrA* codon-optimized for expression in *N. benthamiana*.**

**Figure. S6. Autofluorescence control for GFP signal in agroinfiltrated *N. benthamiana* leaves that do not produce GFP-BfrA**

**Figure S7. SBfrA purification from *N. benthamiana* leaves.**

**Figure S8. *In vitro* nitrogenase activity of reconstituted TSNifH produced in *N. benthamiana* in the presence or absence of BfrA.**

**Table S1. Primers used in this study**

**Table S2. Plasmids used for *N. benthamiana* expression**

## Notes

### Competing Interest Statement

The authors have declared no competing interest.

