## Supplemental Figures for "Enhanced production of nitrogenase components in *Nicotiana benthamiana* through co-expression with Bacterioferritin A"

Fig. S1

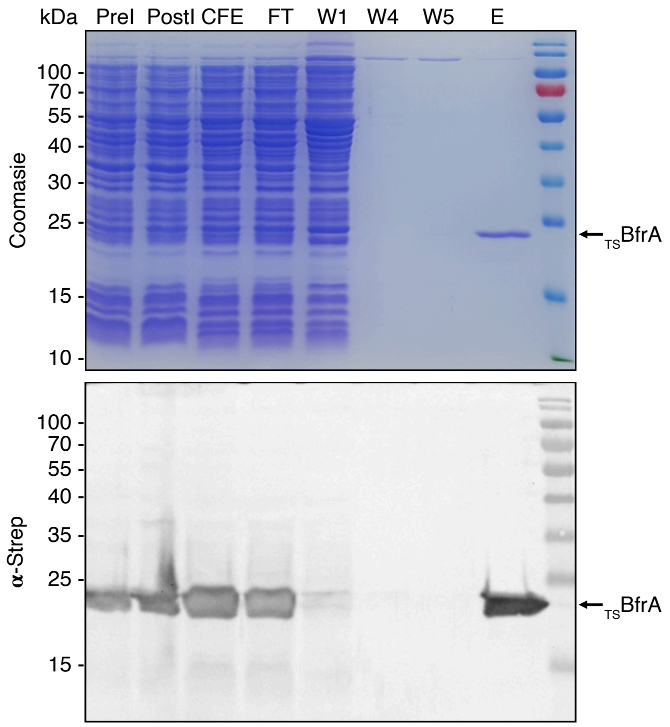


**Figure S1. $_{TS}$BfrA purification from *E. coli*.** Top panel shows Coomassie Brilliant Blue stain of an SDS-PAGE analysis of the purification process of $_{TS}$BfrA from recombinant *E. coli* cells through pre-induction (PreI), post-induction (PostI), cell-free extract (CFE), flowthrough (FT) after passage through a Strep-Tactin column, washes (W1, 4, and 5), and elution (E). Lower panel shows the immunoblot using an anti-Strep antibody of the same samples.

Fig. S2

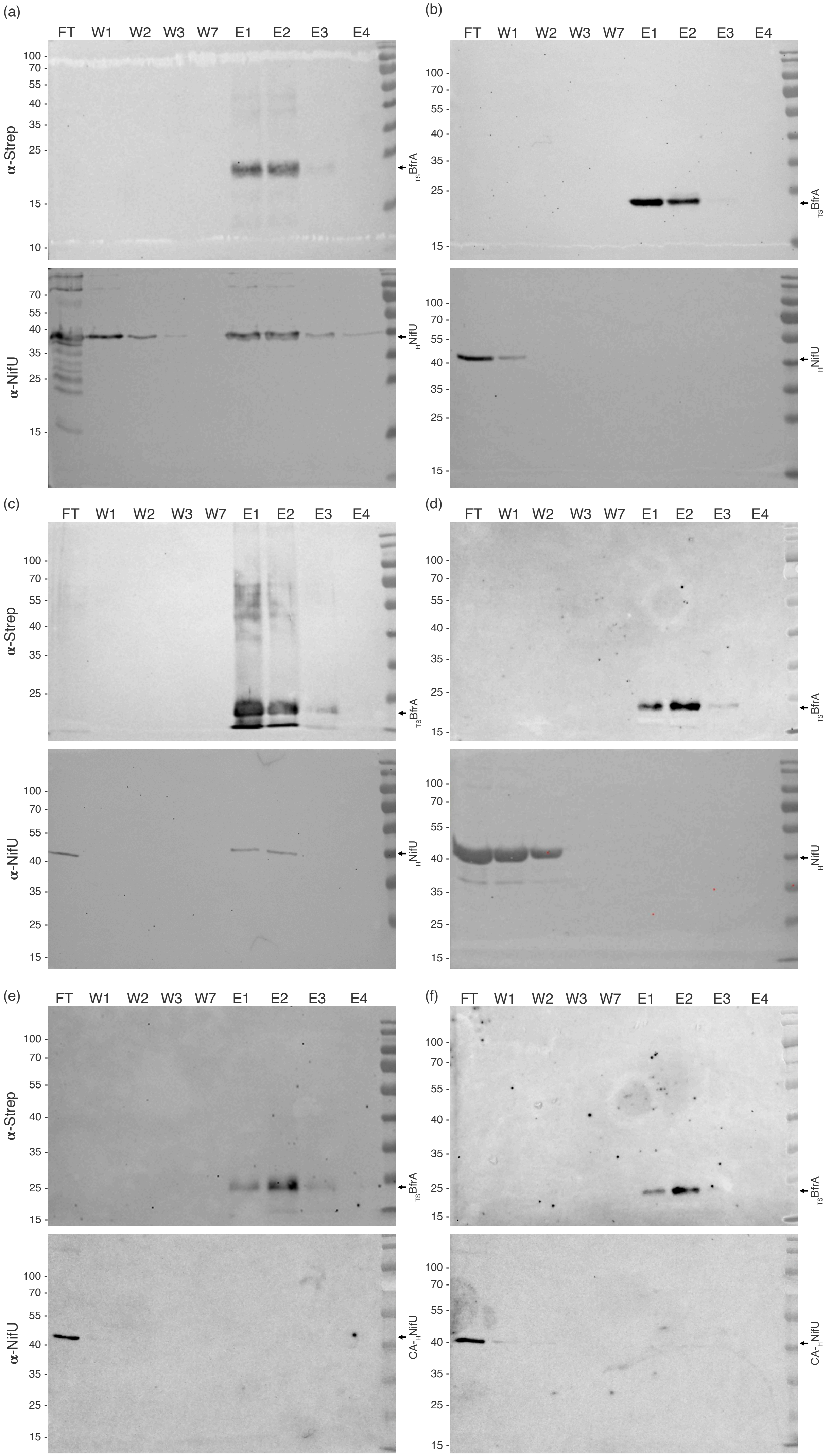


**Figure S2. Uncropped gels of Fig. 1.**

Fig. S3

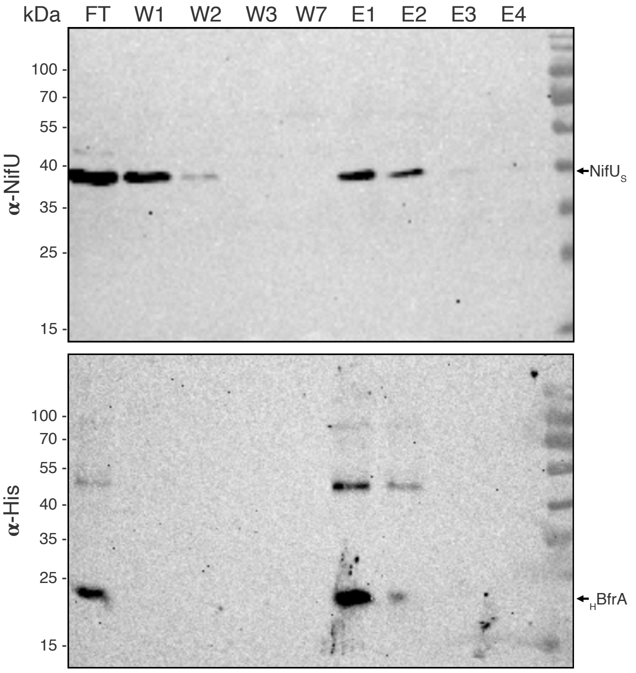


**Figure S3. $NifU_S$ interacts with iron-loaded $_H$BfrA.** Top panel shows the immunodetection with an anti-NifU antibody of Strep-tagged NifU in flowthrough (FT), washes (W1, W3, and W7), and elution (E1, E2, E3, and E4) fractions after being incubated with a histidine-tagged iron-loaded BfrA and passed through a Strep-column. Bottom panel shows the immunoblot of the same fractions developed with an anti-Histidine antibody. Images show a representative assay (n=3).

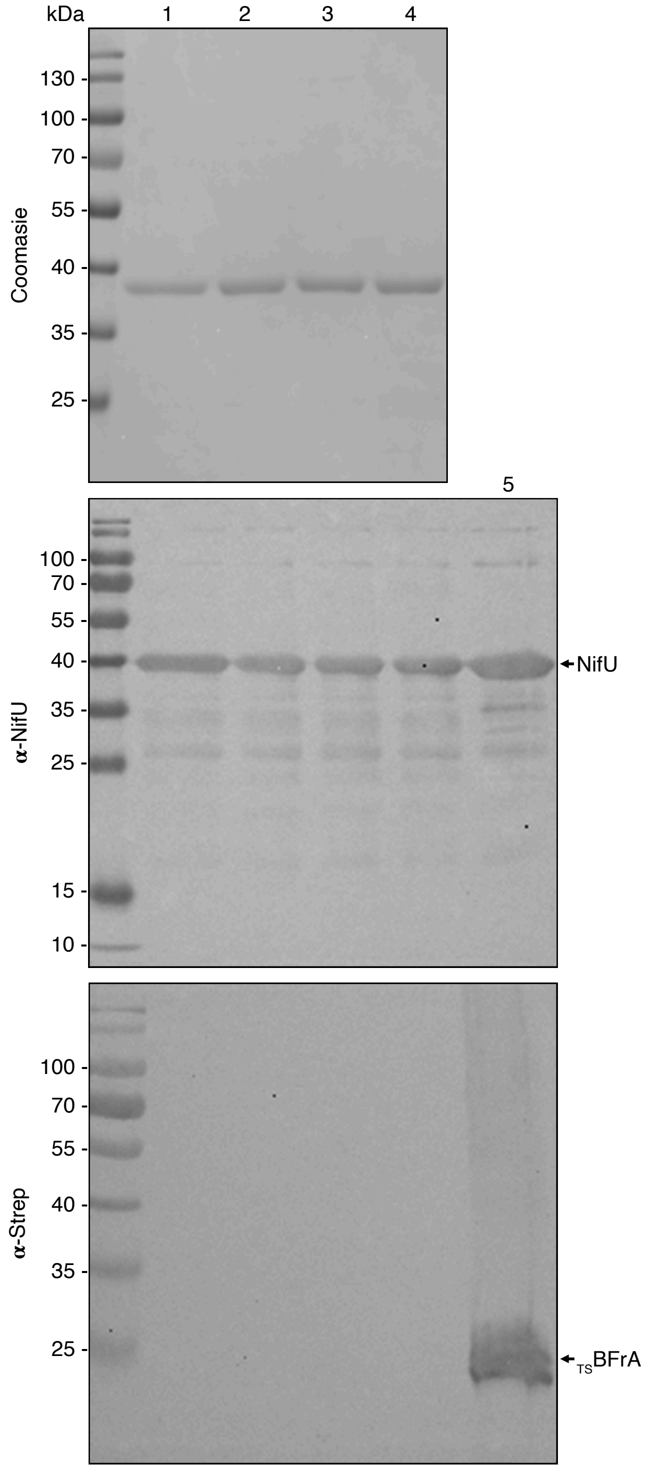


**Figure S4. Control for $_{TS}$BfrA contamination in the samples used for NifU iron content in Fig. 3b.** Top panel shows a Coomassie Brilliant Blue stained SDS-PAGE gel of the flowthrough fractions after passing through a Strep-Tactin column the following samples from Fig. 3b: NifU + $_{TS}$BfrA+TCEP (1), NifU + $_{TS}$BfrA-TCEP (2); the NifU side of the dialysis membrane control (3), and CA-NifU + $_{TS}$BfrA+TCEP (4). Middle panel is the immunoblot of the same fractions using an anti-NifU antibody and pure NifU as a control (5). Bottom panel shows the immunoblot of the same fractions as the top using an anti-Strep antibody and pure $_{TS}$BfrA as control (5).

>*Azotobacter vinelandii* BfrA codon optimized for *Nicotinana benthamiana*

ATGAAGGGAGATAAGATCGTTATCCAACATTTGAATAAGATCCTTGGTAATGA
GTTGATCGCTATTAATCAATATTTTCTTCATGCTAGAATGTACGAAGATTGGGG
ACTTGAGAAGTTGGGTAAACATGAATACCATGAGTCTATCGATGAGATGAAG
CATGCTGATAAGCTTATTAAAAGGATCCTTTTTCTTGAAGGACTTCCAAATTT
GCAAGAGCTTGGAAAGCTTTTGATCGGTGAACATACTAAGGAAATGTTGGAG
TGTGATCTTAAGTTGGAGCAAGCTGGATTGCCTGATCTTAAAGCTGCTATTGC
TTATTGTGAATCTGTTGGAGATTACGCTTCAAGAGAACTTTTGGAGGATATTC
TTGAATCTGAAGAGGATCATATTGATTGGTTGGAGACACAACTTGATTTGATC
GATAAGATCGGTCTTGAAAACTACTTGCAATCTCAGATGGATGAGTAA

**Figure S5. Sequence of *A. vinelandii BfrA* codon-optimized for expression in *N. benthamiana*.**

Fig. S6

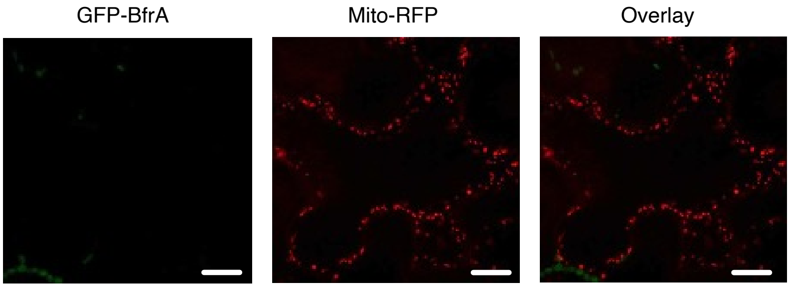


**Figure. S6. Autofluorescence control for GFP signal in agroinfiltrated *N. benthamiana* leaves that do not produce GFP-BfrA.**

Fig. S7

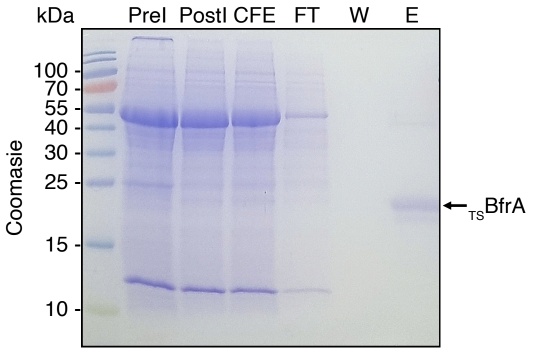


**Figure S7. $_{TS}$BfrA purification from *N. benthamiana* leaves.** Coomassie Brilliant Blue stain of SDS-PAGE gel of the purification process of $_{TS}$BfrA from *N. benthamiana* leaves through pre-inoculation (PreI), post-inoculation (PostI), cell-free extract (CFE), flowthrough (FT) after passage through a Strep-Tactin column, washes (W), and elution (E) fractions.

Fig. S8

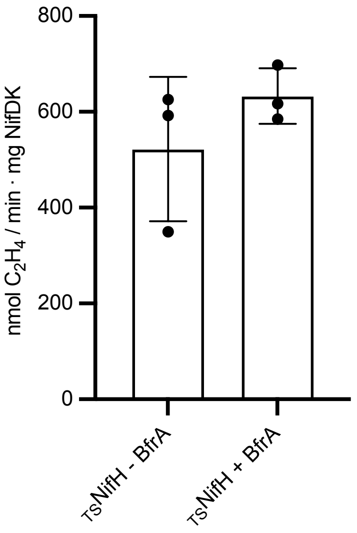


**Figure S8.** ***In vitro*** **nitrogenase activity of reconstituted $_{TS}$NifH produced in *N. benthamiana* in the presence or absence of BfrA.** Bars represent the average ± SD (n=3). * Indicates statistically significant difference ($p$ <0.05).
