## Supplemental Tables for "Enhanced production of nitrogenase components in *Nicotiana benthamiana* through co-expression with Bacterioferritin A"

### Table S1. Primers used in this study

| NAME | SEQUENCE | USE |
|---|---|---|
| Fw eGFP B4 | TGAAGACATCTCAAGCCATGGTG AGCAAGGGCGAG | Prepare GFP with 5´ B4 site for MoClo cloning |
| Rv eGFP B4 | ATGAAGACATCTCGCGAACACTT GTACAGCTCGTCCA | Prepare GFP with 3´ B4 site for MoClo cloning |
| 5´BfrA CONico MoClo B4 | ATGAAGACATCTCAAGCCATGAA GGGAGATAAGATCG | Prepare *N. benthamiana*-codon optimed BfrA with 5´ B4 site for MoClo cloning |
| 5´BfrA CONico MoClo B5 | ATGAAGACATCTCATTCGATGAAG GGAGATAAGATCG | Prepare *N. benthamiana*-codon optimed BfrA with 5´ B5 site for MoClo cloning |
| 3´BfrA CONico MoClo B5 | ATGAAGACATCTCGAAGCTTACTC ATCCATCTGAGATTGC | Prepare *N. benthamiana*-codon optimed BfrA with 3´ B5 site for MoClo cloning |
| 5B*am*HIBfrAinF | ACCATGTTGACCTCCGGATCCGG AGTTGTCGACGAGTCAGTA | In-Fusion cloning |
| 3BamHIBFrAinF | GACAATGCCGAATTCGGATCCAG CGATCTGGATTTTAGTACT | In-Fusion cloning of BfrA |
| 5’ FW 15 nt *Sma*I 20 nt BfrA Up | TAGAGGATCCCCGGGCACGGAAA AAACTGCCGAGT | *bfrA* mutant complementation |
| RV 15 nt *Eco*RI 20 nt Down BfrA | CCATGATTACGAATTCGCGCCGAT TTCGAGCCCAG | *bfrA* mutant complementation |
| *Nde*I_BfrA_prot FW | CCCATATGAAAGGCGATAAGATAG TCATCC | $_{TS}$BfrA cloning |
| *Bam*H1_BfrA_prot_RV | CCGGATCCTTACTCATCCATTTGC GATTGC | $_{TS}$BfrA cloning |
| *Nde*I AvBfrA prot FW | CGTCATATGAAAGGCGATAAGATA GTC | $_{H}$BfrA cloning |
| AvBfrA pET RV | CGGGCTTTGTTAGCAGCCGGATC CTTACTCATCCATTTGCGATTGC | $_{H}$BfrA cloning |

**Table S2. Plasmids used for *N. benthamiana* expression**

| PLASMID | DESCRIPTION | USE | REFERENCE |
|---|---|---|---|
| **pAE160** | COX4 | Mitochondria targeting peptide. Level 0 | Burén *et al*. 2017 |
| **pAE161** | SU9 | Mitochondria targeting peptide. Level 0 | Burén *et al*. 2017 |
| **pAE268** | NifU B5 | *A.vinelandii* NifU codon optimized for *N. benthamiana* cloned in pAGM9121 | Eseverri. *et al* 2020 |
| **pAE33** | NifS B4-B5 | *A.vinelandii* NifS codon optimized for *N. benthamiana* cloned in pAGM9121 | Eseverri *et al*. 2020 |
| **pAE34** | NifU B4-B5 | *A.vinelandii* NifU codon optimized for *N. benthamiana* cloned in pAGM9121 | Eseverri *et al*. 2020 |
| **pAE35** | TS | Twin-Strep Tag codon optimised for *N. benthamiana*. Level 0 | Eseverri *et al*. 2020 |
| **pAE373** | 35Sp (2x; 0,8kb) Ω COX4 TS-NifH TNOS | Mitochondria-targeted $_{TS}$NifH cloned in level 1 position 1 | Jiang *et al*. 2021 |
| **pAE379** | 35SP (Long) SU9 NifM T35S | Mitochondria-targeted NifM cloned in level 1 position 2 | Jiang *et al*. 2021 |
| **pAE738** | AtUbq10m | *Arabidopsis thaliana* Ubiquitin 10 promoter *At4G05320*. Level 0 | Meile *et al*. 2025 |
| **pAGM4673** | MoClo Level 2 acceptor Vector | Level 2 cloning | http://www.addgene.org/48014/ |
| **pAGM8031** | MoClo Level M Acceptor Vector. Position 1 | Assembly of level P components | http://www.addgene.org/48037/ |
| **pAGM8067** | MoClo Level M Acceptor Vector. Position 4 | Assembly of level P components | http://www.addgene.org/48040/ |
| **pAGM9121** | MoClo Level 0 acceptor Vector | Assembly of level 1 components | http://www.addgene.org/51833/ |
| **pAP186** | pICH87633 + pICH44022 + pICH41421 + pICH47751 | P19 silencing suppressor gene cloned in level 1 position 3 | Jiang *et al*. 2021 |
| **pICH41373** | 35Sp (Long) | Promoter (1.3 kb), 35S (Cauliflower Mosaic virus). Level 0 | http://www.addgene.org/50252/ |
| **pICH41414** | 35S Terminator | 3'UTR, polyadenylation signal/terminator, 35S (Cauliflower Mosaic Virus). Level 0 | http://www.addgene.org/50337/ |
| **pICH41421** | NOS Terminator | 3'UTR, Terminator Nos (*A. tumefaciens*). Level 0 | http://www.addgene.org/50339/ |
| **pICH41531** | GFP | CDS Green Fluorescence Protein (*A. victoria*). Level 0 | Eseverri *et al*. 2020 |
| **pICH41744** | End-linker 2. Level 2 | Adapter plasmid used to reconcile overlapping sticky ends between transcriptional units and higher-level destination vector | http://www.addgene.org/48017/ |
| **pICH41766** | End-linker 3. Level 2 | Adapter plasmid used to reconcile overlapping sticky ends between transcriptional units and higher-level destination vector | http://www.addgene.org/48018/ |
| **pICH41780** | End-linker 4. Level 2 | Adapter plasmid used to reconcile overlapping sticky ends between transcriptional units and higher-level destination vector | http://www.addgene.org/48019/ |
| **pICH44022** | P19 | Suppressor of gene silencing (Tomato Bushy Stunt Virus) | http://www.addgene.org/50330/ |
| **pICH47732** | MoClo Level 1 Acceptor Vector Position 1. | Assembly of level 2 components | http://www.addgene.org/48000/ |

| | | | |
|---|---|---|---|
| **pICH47742** | MoClo Level 1 Acceptor Vector Position 2. | Assembly of level 2 components | http://www.addgene.org/48001/ |
| **pICH47751** | MoClo Level 1 Acceptor Vector Position 3. | Assembly of level 2 components | http://www.addgene.org/48002/ |
| **pICH47761** | MoClo Level 1 Acceptor Vector Position 4. | Assembly of e level 2 components | http://www.addgene.org/48003/ |
| **pICH47772** | MoClo Level 1 Acceptor Vector Position 5. | Assembly of level 2 components | http://www.addgene.org/48004/ |
| **pICH47781** | MoClo Level 1 Acceptor Vector Position 6. | Assembly of level 2 components | http://www.addgene.org/48005/ |
| **pICH47791** | MoClo Level 1 Acceptor Vector Position 7. | Assembly of level 2 components | http://www.addgene.org/48000/ |
| **pICH50892** | End-linker 3. Level M | Adapter plasmid used to reconcile overlapping sticky ends between transcriptional units and higher-level destination vector | http://www.addgene.org/48046/ |
| **pICH50927** | End-linker 6. Level M | Adapter plasmid used to reconcile overlapping sticky ends between transcriptional units and higher-level destination vector | http://www.addgene.org/48049/ |
| **pICH51277** | 35Sp (0,4kb) Ω | Promoter (0.4 kb), 35S + 5'UTR, omega (Tobacco Mosaic Virus). Level 0 | http://www.addgene.org/50268/ |
| **pICH51288** | 35Sp (2x; 0,8kb) Ω | Promoter (double), 35S+ 5'UTR, omega. Level 0 | http://www.addgene.org/50269/ |
| **pICH54044** | "Dummy" part level 1 position 4 | Assembly of level 2 or level P components | http://www.addgene.org/48068/ |
| **pICH75322** | MoClo Level P Acceptor Vector | Level P cloning | http://www.addgene.org/48051/ |
| **pICH79300** | End-linker 6. Level P | Adapter plasmid used to reconcile overlapping sticky ends between transcriptional units and higher-level destination vector | http://www.addgene.org/48063/ |
| **pICH87633** | Nosp Ω | Promoter Nos, (*A. tumefaciens*) + 5'UTR, Omega (Tobacco Mosaic Virus). Level 0 | http://www.addgene.org/50271/ |
| **pJQ0016** | pAE738 + pAE161 + pJQ0024 + pICH41414 + pICH47761 | Mitochondria-targeted BfrA cloned in level 1 position 4 | This work |
| **pJQ0018** | pJQ0046 + pJQ0048 + pJQ0047 + pJQ0016 + pICH41766 + pAGM4673 | Level 2 vector targeting $_{TS}$NifU, BfrA, and NifS to mitochondria | This work |
| **pJQ0024** | BfrA B4-B5 | *A. vinelandii* BfrA codon optimized for *N. benthamiana* cloned in pAGM9121 | This work |
| **pJQ0046** | pICH51288 + pAE160+ pAE35 + pAE268 + pICH41421 + pICH47732 | Mitochondria-targeted $_{TS}$NifU cloned in level 1 position 1 | This work |
| **pJQ0047** | pICH51288 + pICH44022 + pICH41414 + pICH47751 | P19 silencing suppressor gene cloned in level 1 position 3 | This work |
| **pJQ0048** | pICH51277 + pAE161 + pAE33 + pICH41421 + pICH47742 | Mitochondria-targeted NifS cloned in level 1 position 2 | This work |
| **pJQ0049** | BfrA B5 | *A.vinelandii* BfrA codon optimized for *N. benthamiana* cloned in pAGM9121 | This work |
| **pJQ0057** | pICH41373 + pAE161 + pVE47 + pJQ0024 + pICH41414 + pICH47732 | Mitochondria-targeted $_{TS}$BfrA cloned in level 1 position 1 | This work |

| | | | |
|---|---|---|---|
| **pJQ0059** | pJQ0046 + pJQ0048 + pJQ0047 + pICH41766 + pAGM4673 | Level 2 vector targeting $_{TS}$NifU and NifS to mitochondria | This work |
| **pJQ0062** | pAE738 + pAE161 + pAE35 + pJQ0024 + pICH41414 + pICH47732 | Mitochondria-targeted GFP-tagged BfrA cloned in level 1 position 1 | This work |
| **pJQ0066** | pJQ0062 + pVE46 + pICH41744 + pAGM4673 | Level 2 vector targeting $_{TS}$BfrA to mitochondria co-expressing the *P19* gene | This work |
| **pVE190** | pICH41373+pAE160+pAE34 +pICH41414 + pICH47781 | Mitochondria-targeted NifU cloned in level 1 position 6 | This work |
| **pVE347** | pICH41373 + pAE161 + pAE33+ pICH41414 + pICH47772 | Mitochondria-targeted NifS cloned in level 1 position 5 | This work |
| **pVE356** | pICH54044 + pVE347 + pVE190 + pICH50927 + pAGM8067 | Level M position 4 vector containing mitochondria-targeted NifS and NifU | This work |
| **pVE358** | pVE360 + pVE356 + pICH79300 + pICH75322 | Level P vector targeting $_{TS}$NifH, NifM, NifS, and NifU to mitochondria | This work |
| **pVE360** | pAE373 + pAE379 + pAP186 + pICH50892 + pAGM8031 | Level M position 1 vector containing mitochondria-targeted NifH, NifM and p19 | This work |
| **pVE363** | pJQ0016 + pVE358 + pICH79300 + pICH75322 | Level P vector targeting $_{TS}$NifH, NifM, BfrA, NifS, and NifU to mitochondria | This work |
| **pVE46** | pICH87633 + pICH44022 +pICH41421 + pICH47742 | P19 silencing suppressor gene cloned in level 1 position 2 | This work |
| **pVE47** | GFP B4 | GFP from pICH41531cloned in B4 position. Level 0 | This work |
